# Sulfotyrosine, an interaction specificity determinant for extracellular protein-protein interactions

**DOI:** 10.1101/2021.10.29.466493

**Authors:** Valley Stewart, Pamela C. Ronald

## Abstract

Tyrosine sulfation, a post-translational modification, can enhance and often determine protein-protein interaction specificity. Sulfotyrosyl residues (sTyr) are formed by tyrosyl-protein sulfotransferase during maturation in the golgi apparatus, and most often occur singly or as a cluster of two or three sTyr within a six-residue span. With both negative charge and aromatic character, sTyr enables numerous atomic contacts as visualized in binding interface structural models, and so there is no discernible binding site consensus. Found exclusively in secreted proteins, sTyr residues occur in four broad sequence contexts. First, a single sTyr residue is critical for diverse high-affinity interactions between peptide hormones and their receptor in both plants and animals. Second, sTyr clusters within structurally flexible anionic segments are essential for a variety of processes including coreceptor binding to the HIV-1 envelope spike protein during virus entry, chemokine interactions with many chemokine receptors, and leukocyte rolling cell adhesion. Third, a subcategory of sTyr clusters occurs in the context of conserved acidic sequences termed hirudin-like motifs that enable several proteins to interact with thrombin, central to normal blood-clotting. Consequently, many proven and potential therapeutic proteins derived from blood-consuming invertebrates depend on sTyr residues for their activity. Fourth, a few proteins that interact with collagen or other proteins contain one or more sTyr residues within an acidic residue array. Refined methods to direct sTyr incorporation in peptides synthesized both in vitro and in vivo, together with continued advances in MS and affinity detection, promise to accelerate discoveries of sTyr occurrence and function.

## Introduction

Post-translational modifications influence protein activity in many ways. Thus, learning how these modifications act singly and in combination is important for understanding protein function (1,2). One post-translational modification increasingly recognized as critical for diverse extracellular interactions in animals, plants and certain bacteria is tyrosine O-sulfation, forming sulfotyrosyl residues (sTyr or Tys). Sulfation is catalyzed during golgi transit by <u>**t**</u>yrosyl-<u>**p**</u>rotein <u>**s**</u>ulfotransferase (TPST), and therefore occurs only in secreted and membrane-spanning proteins (3).

sTyr was discovered nearly 60 years ago (4), but detecting and documenting this modification remains challenging such that the extent of sTyr occurrence has been incompletely defined (5-7). Although the sTyr sulfate linkage is generally stable in weak acid, it is cleaved by the strong acid used in the Edman degradation method used to determine protein sequences (6,8). Compounding the difficulty, routine MS methods do not reliably detect sTyr residues (9), so specialized protocols are being developed to minimize sulfate loss during peptide ionization and fragmentation (7,10-16).

Research on phosphotyrosyl (pTyr) occurrence and function relies in part on a variety of anti-pTyr antibodies, some with broad recognition and others specific for pTyr residues in defined sequence contexts (17). A commercially available anti-sTyr monoclonal antibody binds sTyr residues with high affinity regardless of flanking sequence and discriminates between peptide sTyr and pTyr residues by several hundred-fold (18).

Nevertheless, this reagent has been used sparingly in sTyr research (11). More recently, two antibodies were identified that may discriminate between different chemokine receptor CCR5 sulfoforms (19). It therefore may be feasible to isolate sequence-specific anti-sTyr antibodies to monitor individual sulfation sites.

Systems-level characterization of pTyr residues often employs initial affinity enrichment with a broad-specificity anti-pTyr antibody (20). High-affinity pTyr-binding ***<u>s</u>**rc* homology **<u>2</u>** (SH2) domains (approximately 100 residues) can provide an inexpensive alternative (20). Therefore, SH2 domains with high affinity and specificity for sTyr may offer a useful means for identifying and enriching sTyr-containing proteins (21,22).

Finally, early research on sTyr occurrence and function was constrained by the availability of synthetic sulfopeptides for use in binding assays. No more than two sTyr could be incorporated in a peptide (23), and some studies substituted pTyr for sTyr (24). Today, homogeneous sulfopeptides can be synthesized in vitro with up to three sTyr residues (25-27), and sulfonyl analogs of sTyr provide increased stability (28).

Separately, sulfopeptides with up to five sTyr residues in all combinations have been synthesized in *Escherichia coli* strains that have an expanded genetic code, in which an amber (UAG)-reading tRNA is charged with exogenous sTyr (29-32). A similar system decodes amber as sTyr in mammalian cells (33), providing the possibility for functionally analyzing individual sTyr residues in vivo.

### Why sTyr?

Consider sTyr in contrast to the familiar phosphotyrosyl (pTyr) residue. Sulfation and phosphorylation both add a functional group that is fully ionized at neutral pH and consequently results in increased side-chain polarity (34). However, sTyr makes weaker hydrogen bonds due to its lesser charge (−1 vs. −2) and smaller dipole moment (34).

Functions for pTyr have been studied extensively: its critical role in cellular growth control was discovered in large part from analyzing tumor virus-encoded oncogenic proteins (35). Tyrosyl protein kinase activity serves as “writer” for signaling, whereas separate phosphoprotein phosphatase activity serves as “eraser.” For signal propagation, small pTyr binding domains such as SH2 provide “reader” function for multi-domain output complexes (36,37). By contrast, sTyr is a long-lived post-translational modification (38), and in most cases the sulfate ester likely is stable in physiological conditions (6). Neither “reader” (e.g., portable sTyr-binding domains) nor “eraser” (e.g., sulfoprotein sulfatase) components are known.

The sTyr residue potentiates protein-protein interactions through dual means: the sulfate group, which can make multiple electrostatic interactions to basic Arg or Lys residues in the binding partner; and the Tyr aromatic ring, which engages in both nonpolar and stacking interactions with diverse binding partner residues. Although pTyr can at least partially replace sTyr for certain peptide-peptide interactions (24,39,40), the sTyr sulfate makes distinct ionic contacts (41,42) and therefore in most cases provides a unique in vivo interaction specificity determinant.

sTyr residues are documented in a variety of extracellular proteins (3,9,27,43-46), and currently are known to occur predominately in four broad contexts (**Fig. 1**): (a) in peptide hormones or their receptors, almost all with a single sTyr residue; (b) in conformationally-flexible segments at the amino-termini of cell-surface proteins, most with two or three sTyr clustered within a span of six residues; (c) in conserved sequences (hirudin-like motifs) that interact with thrombin, all with two or three sTyr clustered within a span of six residues; and (d) in tracts of acidic residues at the amino-termini of certain secreted proteins. Thus, sTyr-based interactions usually involve relatively short protein segments, often within large multidomain proteins.

**Fig. 1.**
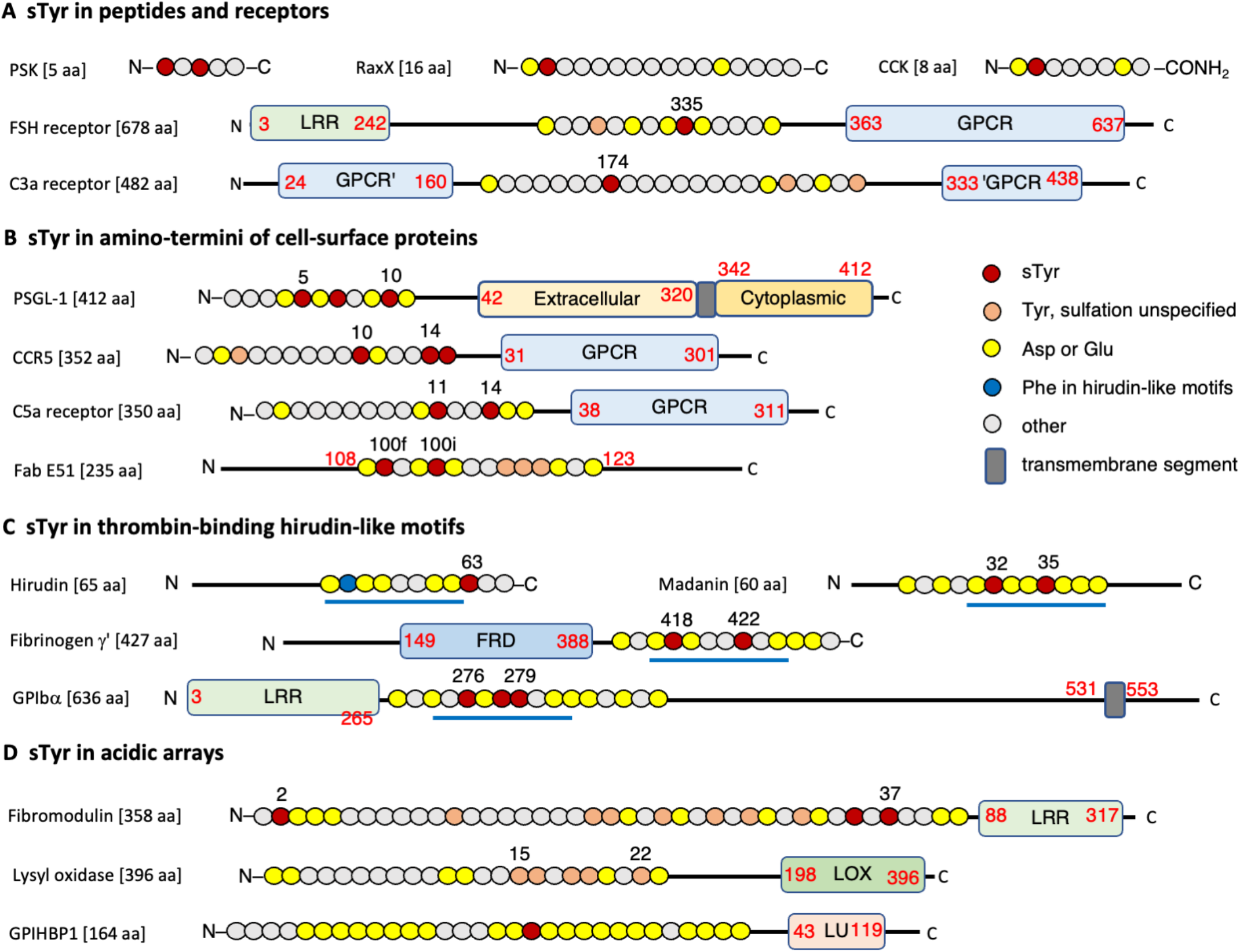
Amino acid sequence contexts for representative sTyr residues. Sketch for journal artist | two-column figure (7.2” X 5.6”) Primary structures of sTyr-containing proteins are shown schematically with major domains drawn approximately to scale. Segments with sTyr residues are depicted as circles, each corresponding to a different amino acid, and colored according to the key. Proteins and peptides are described in the text. The amino-and carboxyl-termini are denoted N and C, respectively, and numbers indicate residue position in the sequence. **A**. sTyr in peptides and receptors. The CCK-8 Phe-amide terminus is denoted CONH_2_. The sTyr-containing sequence in the C3a receptor is in a loop within the seven transmembrane GPCR domain. **B**. sTyr in amino-termini of cell-surface proteins. **C**. sTyr in hirudin-like motifs. The blue lines denote the eight-residue hirudin-like motif. **D**. sTyr in acidic arrays.

Recent discoveries of previously unknown sTyr occurrence (47,48) and functions (26,49-51) show that the repertoire of known sulfoproteins will continue to expand. Here we first describe the enzyme responsible for sTyr formation, and then review examples that illustrate each of these four sTyr contexts (**Fig. 1**). We finish the article by considering some challenges and opportunities for future sTyr research.

### Tyrosyl-protein sulfotransferase (TPST)

Sulfotransferases share similar globular folds that bind the sulfodonor 3’-**<u>p</u>**hospho**<u>a</u>**denosine 5’-**<u>p</u>**hospho**<u>s</u>**ulfate (PAPS) (**Fig. 2A**), the activated intermediate for sulfur assimilation. PAPS synthesis and functions are described elsewhere (52,53).

**Fig. 2.**
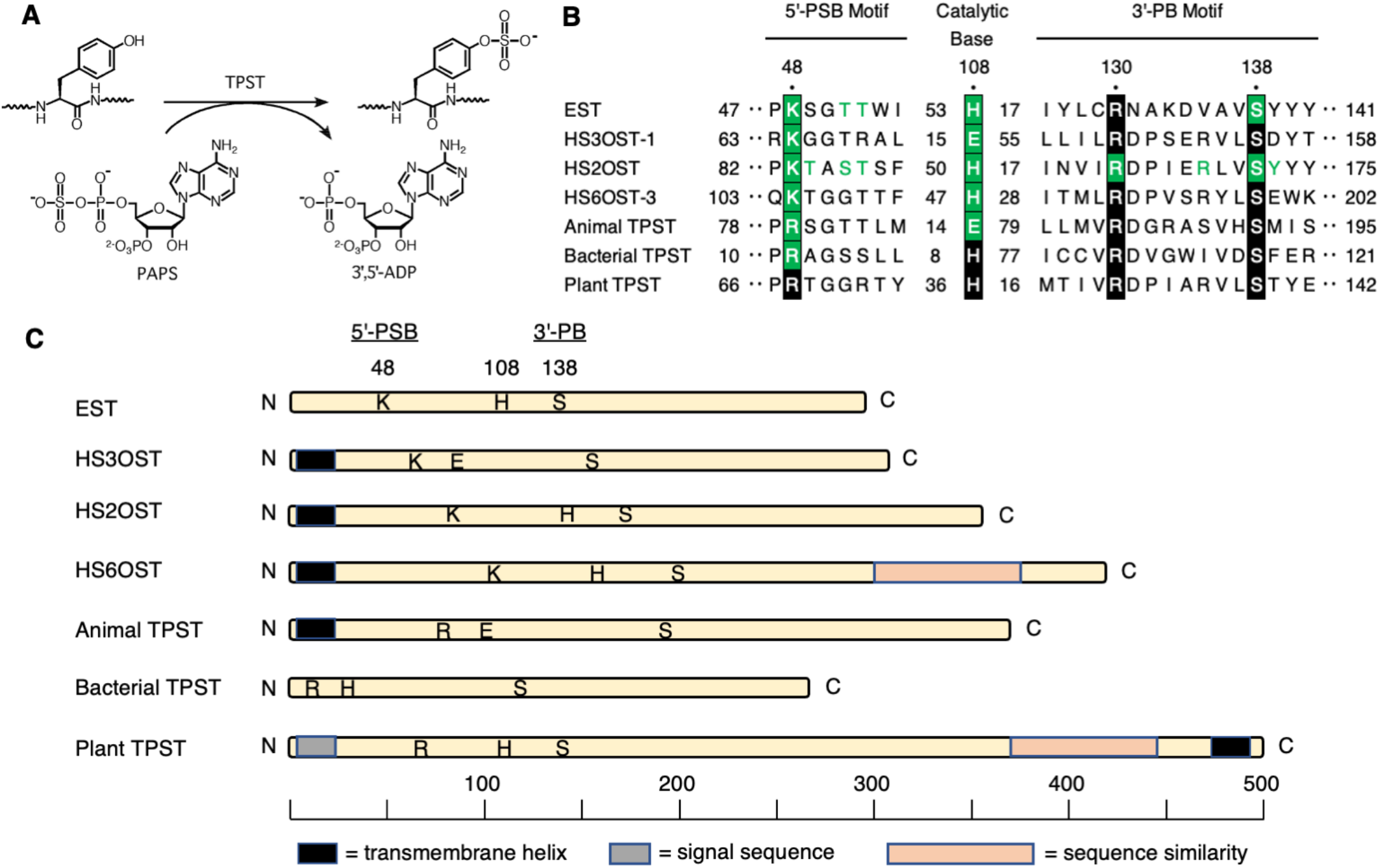
Tyrosyl-protein sulfotransferase (TPST) Sketch for journal artist | two-column figure (7.2” X 4.6”) **A**. Tyrosyl-protein sulfotransferase catalyzes the transfer of sulfate from the universal sulfate donor PAPS (3’-**<u>p</u>**hospho**<u>a</u>**denosine 5’-**<u>p</u>**hospho**<u>s</u>**ulfate) to the hydroxyl group of a peptidyl-tyrosine residue to form a tyrosyl O4-sulfate ester and 3’,5’-ADP (PAP; 3’-**<u>p</u>**hospho**<u>a</u>**denosine 5’-phosphate). Reproduced from reference (5). **B**. Sequence motifs from representative sulfotransferases. Substitutions at residues highlighted in green strongly decrease enzyme activity. Numbers at the left and right ends show positions within the sequence; internal numbers show the number of residues between elements. **C**. Primary structures of representative sulfotransferases, drawn approximately to scale. Letters show the approximate positions of the 5’-PSB (K or R), catalytic base (H or E) and 3’PB (S) motifs shown in panel B. Enzymes are: EST, estrogen 17-β sulfotransferase (*Mus musculus*; PDB code 1AQU); HS3OST-1, heparan sulfate (3-O) sulfotransferase (*M. musculus*; 1S6T); HS2OST, heparan sulfate (2-O) sulfotransferase (*Homo sapiens*; 3F5F); HS6OST-3, heparan sulfate (6-O) sulfotransferase (*Danio rerio*; 5T03); animal TPST (*H. sapiens*; 5WRI); bacterial TPST (*Xanthomonas* spp.; GenBank accession WP_027703307); plant TPST (*Arabidopsis thaliana*; BAI22702).

Two short sequence motifs, termed 5’-PSB and 3’-PB, coordinate the 5’-phosphosulfate and 3’-phosphate groups, respectively (54) (**Fig. 2B**). Sulfotransfer and kinase reactions are similar (**Fig. 2A**), and the geometry of PAPS binding in sulfotransferases is similar to that of ATP binding in nucleotide kinases (55). TPST enzymology has been reviewed elsewhere (56).

Cytoplasmic sulfotransferases termed SULT modify a variety of small molecules such as xenobiotics and hormones, generally share similar overall sequence (57,58). By contrast, tyrosyl-protein and polysaccharide sulfotransferases share little sequence similarity with one another and have different spacing between the 5’-PSB motif, catalytic base residue, and the 3’-PB motif (**Figs. 2B and 2C**), reflecting their engagement with different polymeric substrates.

Metazoan tyrosyl-protein and polysaccharide sulfotransferases are anchored to the Golgi lumen through an amino-terminal transmembrane segment, and thus are type II transmembrane proteins. These sulfotransferases modify proteins and sulfated **<u>g</u>**lycos**<u>a</u>**mino**<u>g</u>**lycans (GAGs) in spatial and temporal coordination with other modifications such as protein glycosylation (2,3,5,43,59,60). Two TPST isoenzymes are encoded by most genera throughout the Metazoa (5). The human TPST isoenzymes are expressed broadly, with one or the other predominating in certain tissues (43,61). Understanding these expression patterns requires defining the relationship between isoenzyme expression and the availability and relative affinity of substrates.

For example, homozygous *tpst-1* null mice have lower average body weight, which may result in part from reduced levels of the sulfated hormones **<u>c</u>**hole**<u>c</u>**ysto**<u>k</u>**inin (CCK) and gastrin (described below). By contrast, homozygous *tpst-2* null mice display primary hypothyroidism, likely resulting from failure to sulfate the receptor for thyroid stimulating hormone (see below). Homozygous *tpst-2* null males also are infertile. Finally, homozygous *tpst-1 tpst-2* double null mice usually die soon after birth due to cardiopulmonary insufficiency (62-66). Thus, TPST-1 and TPST-2 functions overlap only partially. The challenge remains to identify specific bases for most of these phenotypes.

Metazoan TPST displays broad substrate specificity. Most substrates with at least moderate affinity have an Asp residue just proximal to the sulfoaccepting Tyr residue (61,67-71) (**Fig. 1**). Non-conserved flanking residues, often rich in acidic Asp and Glu residues, make multiple interactions with different TPST residues near the active site (67,68,70). Overall, these broader sequence features and their roles in metazoan TPST binding affinity and sulfation efficiency remain obscure (72).

Plants encode a single TPST (53). In the model plant *Arabidopsis thaliana*, a null mutant displays developmental phenotypes such as dwarfism (73), consistent with the functions for the only known plant TPST substrates, sulfopeptide hormones involved in multicellular development (described below).

Plant TPST has a carboxyl-terminal transmembrane segment (type I transmembrane protein) in contrast to metazoan TPST and GAG sulfotransferases (73,74) (**Fig. 2C**). Sequences for plant and metazoan TPST are not obviously related, although plant TPST does share carboxyl-terminal sequence similarity with **<u>h</u>**eparan **<u>s</u>**ulfate **<u>6</u>**-**<u>O</u>**-sulfotransferase (HS6OST), a GAG sulfotransferase (73). This carboxyl terminus is a prominent α-helix in the HS6OST-3 X-ray structure (75) and is not present in other characterized sulfotransferases.

Initial analysis did not detect plant TPST sequence similarity to the conserved 5’-PSB and 3’-PB motifs (73). With more sequences now available, it is evident that different polymer sulfotransferases have PAPS-binding 5’-PSB and 3’-PB motifs with varied sequences and spacings, and so plant TPST sequences do include putative PAPS-binding motifs that match well with those from other polymer sulfotransferases (**Fig. 2B**). Experimental work is necessary to test key residues in the plant TPST putative 5’-PSB and 3’-PB motifs and to examine substrate specificity determinants.

Plant TPST orthologs are conserved throughout green plants including unicellular algae like *Chlamydomonas* spp. that are not known to synthesize sulfopeptide hormones (53,73). It will be interesting to learn the algal substrate proteins for TPST, and to determine if any are conserved in land plants.

Most bacteria, archaea and non-photosynthetic eukaryotic microbes do not encode TPST. Nevertheless, TPST is made by some species of plant-pathogenic bacteria in the genus *Xanthomonas* that also synthesize the RaxX protein (**<u>r</u>**equired for **<u>a</u>**ctivation of Xa21-mediated immunity), a molecular mimic of the PSY sulfopeptide hormone (76,77) (**Fig. 3A**). The bacterial TPST sequence is most similar to that of the Golgi-localized metazoan TPST, except that bacterial TPST acts in the cytoplasm prior to substrate secretion and therefore is not membrane-anchored (78).

**Fig. 3.**
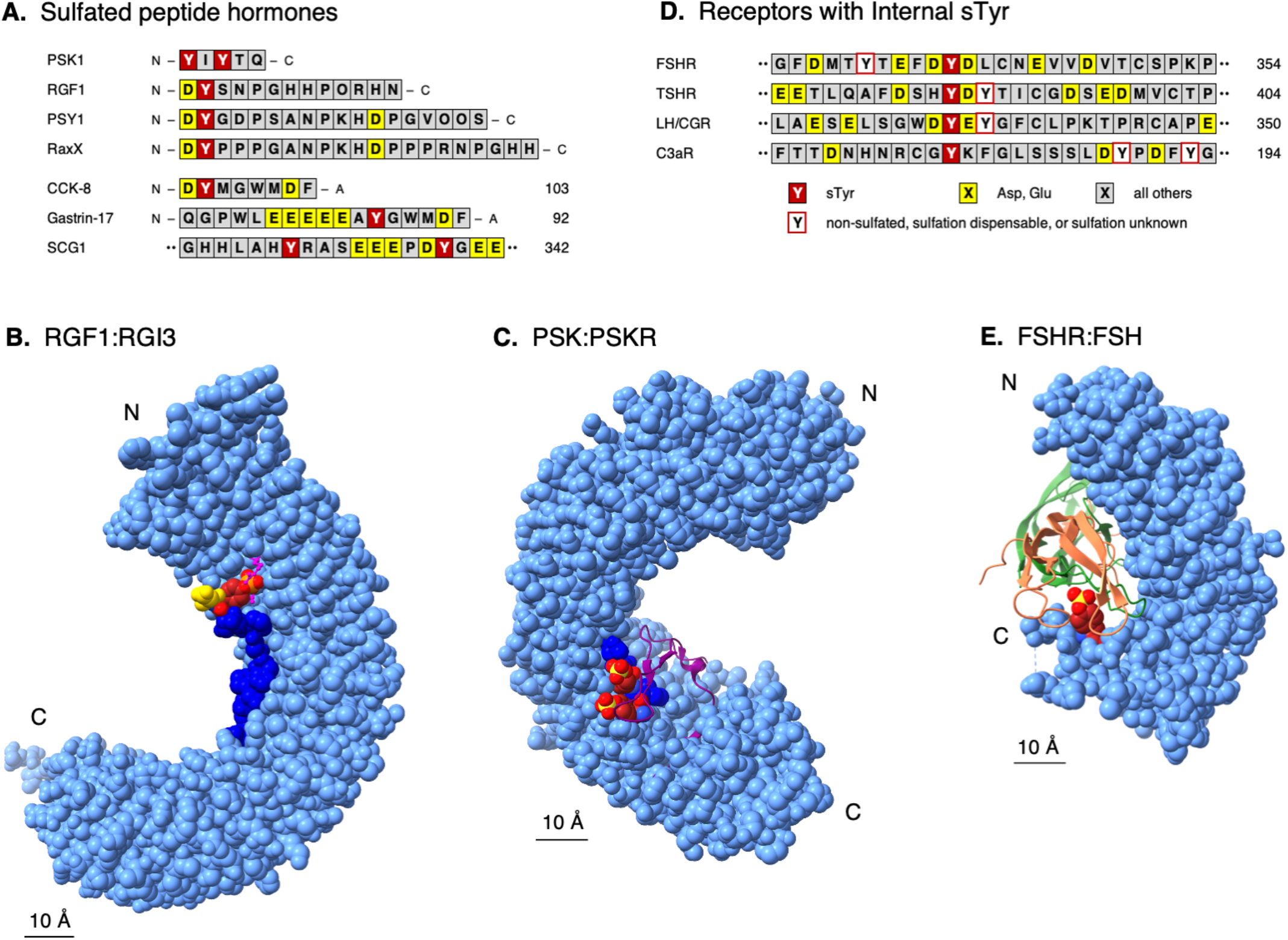
sTyr in peptides and receptors. two-column figure (7.2″ X 5.3″) **A**. sTyr residue contexts are shown for the mature forms of the plant sulfopeptide hormones PSK1 (phytosulfokine), RGF1 (root meristem growth factor), PSY1 (peptide sulfated on tyrosine), the hormone mimic RaxX (required for activating XA21 protein X), and the animal sulfopeptide hormones CCK-8 (cholecystokinin), gastrin-17, and one of two sulfopeptides derived from the secretogranin-1 (SGC-1) precursor. Residues are colored according to the key in panel D. O denotes hydroxyproline. **B**. The co-crystal X-ray model of sulfated RGF (dark blue) bound to the RGI3 (RGF-insensitive-3) receptor ectodomain (light blue) is depicted at 2.6 Å resolution (86) (PDB code 5HYX). Atoms are shown as Van der Waals spheres. The RGI3 ectodomain is built from 23 LRRs (leucine-rich repeats) stacked one upon the next. RGF residue sTyr-2 (dark red; S atom yellow) contacts RGI3 residues Arg-195 and Ala-222, depicted as balls-and-sticks (magenta). RGF residue Asp-1 (yellow) is mostly exposed. **C**. The co-crystal X-ray model of sulfated PSK (dark blue) bound to the PSK receptor ectodomain (light blue) is depicted at 2.5 Å resolution (88) (PDB code 4Z63). Atoms are shown as Van der Waals spheres. The RGI3 ectodomain is built from 21 LRRs (leucine-rich repeats) stacked one upon the next, with a 36-residue island domain, shown as a ribbon (magenta), inserted into LRR number 18. PSK residues sTyr-1 and sTyr-3 (dark red; S atoms yellow) mostly contact residues in the island domain. **D**. sTyr residue contexts are shown for internal segments (**Fig. 1A**) of the receptors for FSH (follicle-stimulating hormone), TSH (thyroid stimulating hormone), LH/CGR (luteinizing hormone/choriogonadotropin) and complement component C3a. **E**. The co-crystal X-ray model of FSH (follicle-stimulating hormone; α subunit salmon; β subunit light green) bound to the sulfated FSH receptor ectodomain (light blue) is depicted at 2.5 Å resolution (93) (PDB code 4AY9). The FSHR ectodomain is built from 12 LRRs stacked one upon the next. FSHR residue sTyr-335 (dark red; S atom yellow) contacts several residues each in both FSH subunits.

### sTyr increases binding affinity for peptide hormone-receptor pairs

Secreted sulfopeptide hormones of roughly 10-20 residues control growth and development in land plants (38,79-81), and digestion, neurotransmission and endocrine functions in animals (82,83). Most plant sulfopeptide hormones possess the invariant amino-terminal residue pair Asp-1 sTyr-2, where the Asp residue presumably directs plant TPST to the substrate Tyr residue (**Fig. 3A**).

Plant peptide hormones bind their cognate receptors through **l**eucine-rich repeat (LRR) extracellular domains (84,85). The plant sulfopeptide hormones tested display high affinities for their cognate receptor LRR domains, ranging from 1 to about 300 nM, and in each case the sTyr residue contributes tenfold or more to affinity compared with the nonsulfated peptide. (48,78,86,87).

Plant receptor binding to the sTyr residue is shown in X-ray co-crystal structures for the sulfopeptide hormones **<u>r</u>**oot meristem **<u>g</u>**rowth **<u>f</u>**actor (RGF) and **<u>C</u>**asparian strip **<u>i</u>**ntegrity factor (CIF) with their cognate receptor LRR domains (86,87). Both structures reveal overall hydrophobic sTyr-binding interfaces, but only one Arg residue is conserved among residues that make specific contacts (**Fig. 3B**). Based on these two examples, it appears that plant LRR receptor sTyr binding interfaces have few sequence constraints and therefore might readily be formed during evolution. Further examples of LRR-sulfopeptide interactions are needed to determine the limits of these constraints.

The pentapeptide **<u>p</u>**hyto**<u>s</u>**ulfo**<u>k</u>**ine (PSK) is the only known plant hormone that contains two sTyr residues, which together confer about 40-fold higher affinity for the PSK receptor (38,88,89). The PSK receptor LRR domain contains a 36 residue “island” that contacts both sTyr residues through multiple interactions (88) (**Fig. 3C**). These different features illustrate how sTyr residues are versatile interaction determinants even in a broadly similar context such as sulfopeptide-LRR binding.

The animal sulfopeptide hormones CCK and gastrin are processed to generate different length bioactive peptides with a common carboxyl-terminal Phe-amide (**Fig. 3A**). The CCK sTyr residue mostly is sulfated (83), whereas the gastrin sTyr residue is heterogeneously sulfated, due perhaps to a low-affinity sequence environment for TPST recognition (67).

CCK and gastrin signal through the homologous G protein-coupled receptors CCK1R and CCK2R. CCK1R binds sulfated CCK with up to one-thousand-fold greater affinity than nonsulfated CCK, sulfated gastrin, or nonsulfated gastrin. In contrast, the CCK2R receptor makes little distinction (90). Thus, ligand sulfation state is a specificity determinant for CCK1R-CCK interaction. No X-ray structure is available for either CCK1R or CCK2R, but various studies indicate that these sulfopeptides interact across a broad region of the receptor external face (82), and that a conserved Arg residue in a CCK1R extracellular loop interacts specifically with the CCK sTyr (91).

Mirroring these interactions between nonsulfated receptors and sulfated peptide hormones, some nonsulfated ligands bind single sTyr-containing receptors (**Fig. 3D**). Glycoprotein hormones such as follicle-stimulating hormone are central to the complex endocrine system that regulates normal growth, sexual development, and reproductive function. Receptors for glycoprotein hormones comprise an amino-terminal LRR domain connected through a flexible region to a carboxyl-terminal G protein-coupled receptor (**Fig. 2A**). This interdomain region contains an sTyr residue that is indispensable for hormone recognition and signaling (92,93).

Upon binding to the LRR domain, follicle-stimulating hormone (about 200 residues) exposes a hydrophobic interface that contains two positively charged residues for electrostatic interactions with the sTyr sulfate (92,93) (**Fig. 3E**). Homologous receptors for thyroid stimulating hormone and leutenizing hormone similarly contain an essential interdomain flexible region with a single sTyr residue (94) (**Fig. 3D**).

Component C3a, a 77 residue peptide generated through the complement cascade, acts through its receptor to stimulate many aspects of the inflammatory response, sometimes leading to anaphylaxis (95). The G protein-coupled C3a receptor has an sTyr residue, essential for binding C3a, within an unusually large extracellular loop (96) (**Fig. 3D**). To date, an atomic-level view of receptor binding and activation is not available (97).

### sTyr clusters mediate a variety of protein-protein interactions

In most cases sTyr residues occur in clusters of two or three within a six-residue span. This arrangement enables a wide range of interactions as illustrated by the examples below.

### sTyr clusters are essential for HIV-1 entry

HIV-1 binding and entry depends on the viral envelope **<u>g</u>**lyco**<u>p</u>**rotein (gp) spike, a trimer of gp120-gp41 heterodimers (98). First, the spike binds to the cell-surface receptor CD4, thereby altering gp120 conformation to expose its bridging sheet element (99). Then, the coreceptor amino terminus binds the bridging sheet, triggering membrane fusion and viral entry (98,100,101).

The principal T cell coreceptors for HIV-1 entry are the chemokine receptors CCR5 and CXCR4 (102). These are G protein-coupled receptors for **<u>chemo</u>**tactic cyto**<u>kines</u>** (chemokines), secreted signaling proteins of approximately 75 residues that direct leukocyte movement toward injury or infection sites (103). The amino-terminal segment in many chemokine receptors includes an sTyr cluster that is essential for high-affinity chemokine binding (44,46,104-107) (**Fig. 2B & Fig. 4A**). Interest in sTyr function accelerated with discoveries that the CCR5 sTyr cluster is essential for HIV-1 entry (100) and is mimicked in some HIV-1 broadly-neutralizing antibodies (108).

**Fig. 4.**
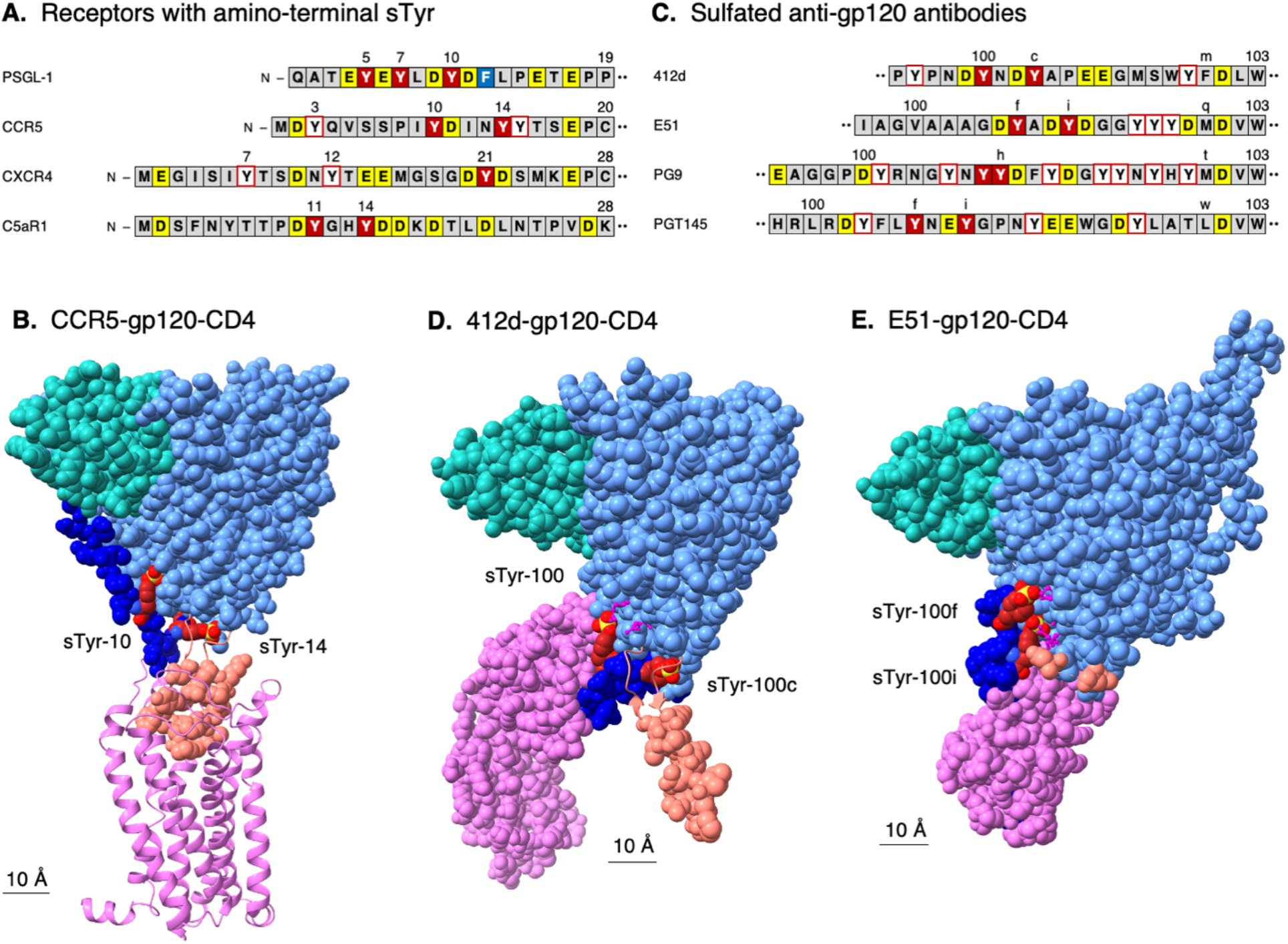
sTyr in receptor amino-termini and in anti-gp120 antibodies. two-column figure (7.2” X 5.3”) **A**. sTyr cluster contexts are shown for the amino-termini of the cell-surface proteins PSGL-1 (P-selectin glycoprotein ligand-1), CCR5 (CC-type chemokine receptor 5), CXCR4 (CXC-type chemokine receptor 4) and C5aR (complement component C5a receptor). Residues are colored according to the key in **Fig. 3D**. **B**. The cryo-EM model of HIV-1 gp120 (glycoprotein 120; light blue) bound to host CCR5 (light purple) and one subdomain of host CD4 (cluster of differentiation 4 receptor; light green) is depicted at 3.9 Å resolution (101) (PDB code 6MEO). Atoms are shown as Van der Waals spheres except for CCR5, depicted as ribbons the better to display the seven transmembrane helix structure and to visualize interaction with the gp120 V3 loop residues (salmon). The CCR5 amino-terminal segment (dark blue) includes residues sTyr-10 and sTyr-14 (dark red; S atoms yellow) that interact with residues in the gp120 bridging sheet element including the base of the V3 loop. Certain residues at the base of the V3 loop are shown as ribbons the better to visualize the sTyr residues. **C**. sTyr cluster contexts are shown for the HCD3 regions from representative CD4i anti-gp120 antibodies: 412d (PDB code 2QAD); E51 (6U0L); PG9 (3U2S); PGT145 (3U1S). **D**. The co-crystal X-ray model of HIV-1 gp120 (glycoprotein 120; light blue) bound to the antibody 412d heavy chain (light purple) and one subdomain of host CD4 (light green) is depicted at 3.3 Å resolution (112) (PDB code 2QAD). Certain residues at the base of the V3 loop are shown as ribbons the better to visualize the sTyr residues. The HCD3 region (dark blue) includes residues sTyr-100 and sTyr-100c (dark red; S atoms yellow) that interact with residues in the gp120 bridging sheet element including the base of the V3 loop. Certain residues at the base of the V3 loop are shown as ribbons the better to visualize the sTyr residues. Only a portion of the 412d light chain is shown. **E**. The cryo-EM model of HIV-1 gp120 (light blue) bound to the antibody E51 heavy chain (light purple) and one subdomain of host CD4 (light green) is depicted at 3.3 Å resolution (117) (PDB code 6U0L). The HCD3 region (dark blue) includes residues sTyr-100f and sTyr-100i (dark red; S atoms yellow) that interact with residues in the gp120 bridging sheet element. The V3 loop is not resolved in this structure; gp120 residues at the V3 base (salmon) indicate its position. Only a portion of the E51 light chain is shown.

CCR5 residues sTyr-10 and sTyr-14 are necessary for proper gp120-CCR5 interaction, whereas sulfation at residues Tyr-3 or Tyr-15 dispensable (109-111) (**Fig. 4A**). A cryo-EM structural model shows the CCR5 amino-terminal segment in an extended conformation across the gp120 bridging sheet. In this model, residue sTyr-14 is mostly buried in a pocket lined with hydrophobic contacts and capped by electrostatic interactions. By contrast, residue sTyr-10 is mostly exposed (98,101) (**Fig. 4B**).

Certain HIV-1-infected individuals produce antibodies that recognize the **<u>CD4</u>**-**<u>i</u>**nduced (CD4i) conformation of gp120 with the bridging sheet exposed (108,112). Most of these CD4i antibodies contain **<u>h</u>**eavy chains with unusually long and diverse sequences for the third **<u>c</u>**omplementarity-**<u>d</u>**etermining regions (**HCD3**) that form protruding loops to contact otherwise poorly accessible epitopes (112-114). Many of these CD4i antibody HCD3 sequences include sTyr residues required for gp120 binding (108,112) (**Fig. 4C**).

In an X-ray co-crystal structure with the gp120-CD4 complex, residues sTyr-100c^412d^ and sTyr-100^412d^ bind the same gp120 interfaces as the modeled residues sTyr-10^CCR5^ and sTyr-14^CCR5^, respectively (101,115) (**Fig. 4D**). (The antibody numbering accounts for variable numbers of residues, labeled 100a-100w, between HCD3 residues at positions 100 and 101.) Separate studies with CCR5 sulfopeptides and antibody 412d sulfoforms expressed with the expanded genetic code reveal that the the buried residues sTyr-14^CCR5^ and sTyr-100c^412d^ are essential for binding, whereas the more exposed residues sTyr-10^CCR5^ and sTyr-100^412d^ are ancillary (111,116). Thus, the CCR5 and 412d sTyr-containing elements apparently share similar interaction with gp120.

A cryo-EM structural model shows a different binding pattern by antibody E51 (117) (**Fig. 4E**). Although residue sTyr-100i^E51^ binds to the same gp120 interface as the exposed ancillary residues sTyr-10^CCR5^ and 412d sTyr-100^412d^, residue sTyr-100f^E51^ instead makes unique interactions (**Fig. 4E**). Interestingly, for antibody E51 sulfoforms, exposed residue sTyr-100i^E51^ is essential whereas the unique residue sTyr-100f^E51^ is ancillary (32).

Thus, antibody E51 sTyr-gp120 interactions are distinct from those for chemokine receptor CCR5 and antibody 412d. Intriguingly, E51 also is more potent than 412d (117), and forms the basis for an effective anti-HIV peptide (118). Apparently the gp120 bridging sheet is versatile enough to make high-affinity interactions with different sTyr-containing flexible segments, increasing opportunities to meet the stereochemical constraints to binding sulfate (112).

### sTyr clusters contribute to combinatorial receptor-peptide interactions

Human cells express about 50 chemokines and about 20 chemokine receptors acting in different combinations in different tissues (119), so binding interface versatility is a hallmark. Roughly half of the chemokine receptors contain sTyr residues in their amino-termini (44). The sTyr cluster forms the core for binding the chemokine ligand globular domain and helps enable a given receptor to interact with multiple chemokines (46,104,107,119). The CCR5 amino-terminal segment has yet to be captured in a receptor-chemokine X-ray or cryo-EM structure (44,101,107,120,121), suggesting that it is intrinsically disordered (122).

Indeed, NMR spectroscopy and modeling suggests that CCR5-CCL5 interactions are highly dynamic (121,123). In accordance with these observations, separate sulfopeptide binding experiments revealed that any combination of two sTyr residues at CCR5 positions 10, 14 and 15 enables strong binding to chemokine CCL5 (124).

Together, these results support the hypothesis that the CCR5 flexible sTyr-containing anionic segment makes a variety of dynamic yet high-affinity contacts to the chemokine, in striking contrast to the relatively fixed contacts observed in the CCR5-gp120 interactions described above (121,124).

Chemokines also bind sulfated GAGs to form chemotactic gradients (103). Notably, GAG sulfate groups make electrostatic interactions with many of the same chemokine basic residues that contact sTyr sulfates. Thus, chemokine functionality is expanded by a single versatile interface that binds different sulfated polymers for different purposes (106,123,125).

Chemokines provide signals for leukocyte movement toward sites of infection or injury. This rolling cell adhesion is mediated in part by **<u>P</u>**-**<u>s</u>**electin **<u>g</u>**lycoprotein **<u>l</u>**igand-**<u>1</u>** (PSGL-1), a homodimeric mucin-like cell surface glycoprotein. PSGL-1 contains three sTyr residues in its amino-terminal flexible segment (**Fig. 4A**) and engages with the cell-surface adhesion P-selectin through a mechanosensitive catch-bond that enables rapid engagement and release under force in the bloodstream flow (126,127). An X-ray co-crystal structure of a short PSGL-1 sulfoglycopeptide bound to the P-selectin amino-terminal domain shows residue sTyr-7 with several electrostatic and hydrophobic contacts to residues in P-selectin, and residue sTyr-10 with multiple hydrophobic contacts that orient the sulfate for hydrogen bonding to a critical Arg residue (2,128).

Although all three PSGL-1 sTyr residues are necessary for full P-selectin interactions, any one sTyr suffices for partial function in a variety of assays relevant to rolling cell adhesion (129). Indeed, residue sTyr-5 is not visible in the P-selectin-PSGL-1 cocrystal structure, suggesting that the observed structure may represent one of several productive conformations (128). In this hypothesis, the sTyr cluster potentially presents a variety of conformations suitable for interaction. Nevertheless, a separate study identified sTyr-7 as essential for PSGL-1 binding to P-selectin (130), congruent with the extensive interactions made by this residue (128).

Separately, PSGL-1 helps regulate aspects of T cell function, including the progressive loss of effector function in so-called exhausted T cells that accompanies persistent antigen stimulation (131). In this context, PSGL-1 is the receptor for **<u>V</u>**-domain **<u>i</u>**mmunoglobulin **<u>s</u>**uppressor of **<u>T</u>** cell **<u>a</u>**ctivation (VISTA), a pH-responsive T cell inhibitor (131). In a computational model, all three PGSL-1 sTyr residues make ionic interactions with His residues along one edge of VISTA. In this model, sTyr interaction depends upon His protonation, as would occur in acidic tumor microenvironments (pH ≤ 6) (49). Sulfation is critical for VISTA-PSGL-1 binding, but contributions of individual sTyr residues are not known (49). Together, interactions with P-selectin and VISTA illustrate how sTyr cluster structural flexibility enables impressive functional versatility by the PSGL-1 amino-terminal segment.

Complement protein C5a (74 residues), generated through the complement cascade, stimulates inflammatory mediator release and also is a potent chemoattractant (95). Similar to chemokine receptors, the G protein-coupled C5a receptor amino-terminal segment contains two sTyr residues (132) (**Fig. 4A**). **<u>Ch</u>**emotaxis **<u>i</u>**nhibitory **<u>p</u>**rotein of **<u>*S*</u>***taphylococcus aureus* (CHIPS; 121 residues) also binds the amino-terminal segment, thereby inhibiting C5a-dependent inflammatory responses (97). The C5a receptor sTyr residue pair increases CHIPS binding affinity for peptides from the C5a receptor amino-terminal segment by approximately 1,000-fold (133). In the NMR structure of CHIPS bound to the C5a receptor amino-terminal peptide, the two sTyr residues are within a five residue β-strand and make several hydrophobic and electrostatic contacts to residues in CHIPS (133).

These three examples – CCR5, PSGL-1, and C5aR – illustrate a variety of sTyr interactions ranging from relatively ordered, as with the sulfoantibody-gp120 and C5aR-CHIPS interactions, to highly dynamic, as with the CCR5-CCL5 and PSGL-1-P-selectin interactions. Thus, it appears that two sTyr residues can enable a wider range of binding modes and partners than observed in the single sTyr residue hormones and receptors described above.

### sTyr clusters in hirudin-like motifs bind the blood clotting enzyme thrombin

The sTyr cluster-containing anionic segments described above are versatile and dynamic, and there is no obviously conserved sequence pattern. Here, a different collection of anionic segments makes conserved interactions and contain conserved sTyr cluster sequences. Thus, sTyr clusters participate across a range of anionic segment functions.

Aberrant hemostasis – ranging from profuse bleeding to excessive clotting – underlies or complicates several human conditions, and many treatments have been developed (134-136). The serine endoprotease α-thrombin is central to controlling the balance between initiating and terminating blood clotting (thrombosis). Thrombin function depends on its interactions with molecular tethers in several different substrates and inhibitors (137,138). This number and diversity of thrombin binding partners provides an opportunity to explore unity and diversity in sTyr cluster interactions.

One clot-inhibiting therapeutic used for over one hundred years is hirudin, an sTyr-containing **<u>d</u>**irect **<u>t</u>**hrombin **<u>i</u>**nhibitor (DTI) made by *Hirudo* leeches (139,140). Hirudin (65 residues) comprises a carboxyl-terminal anionic segment that binds thrombin and an amino-terminal domain that occludes the thrombin active site. Similar anionic segments, termed hirudin-like motifs (141), are present in other thrombin-binding proteins (137,138) (**Fig. 5A**).

**Fig. 5.**
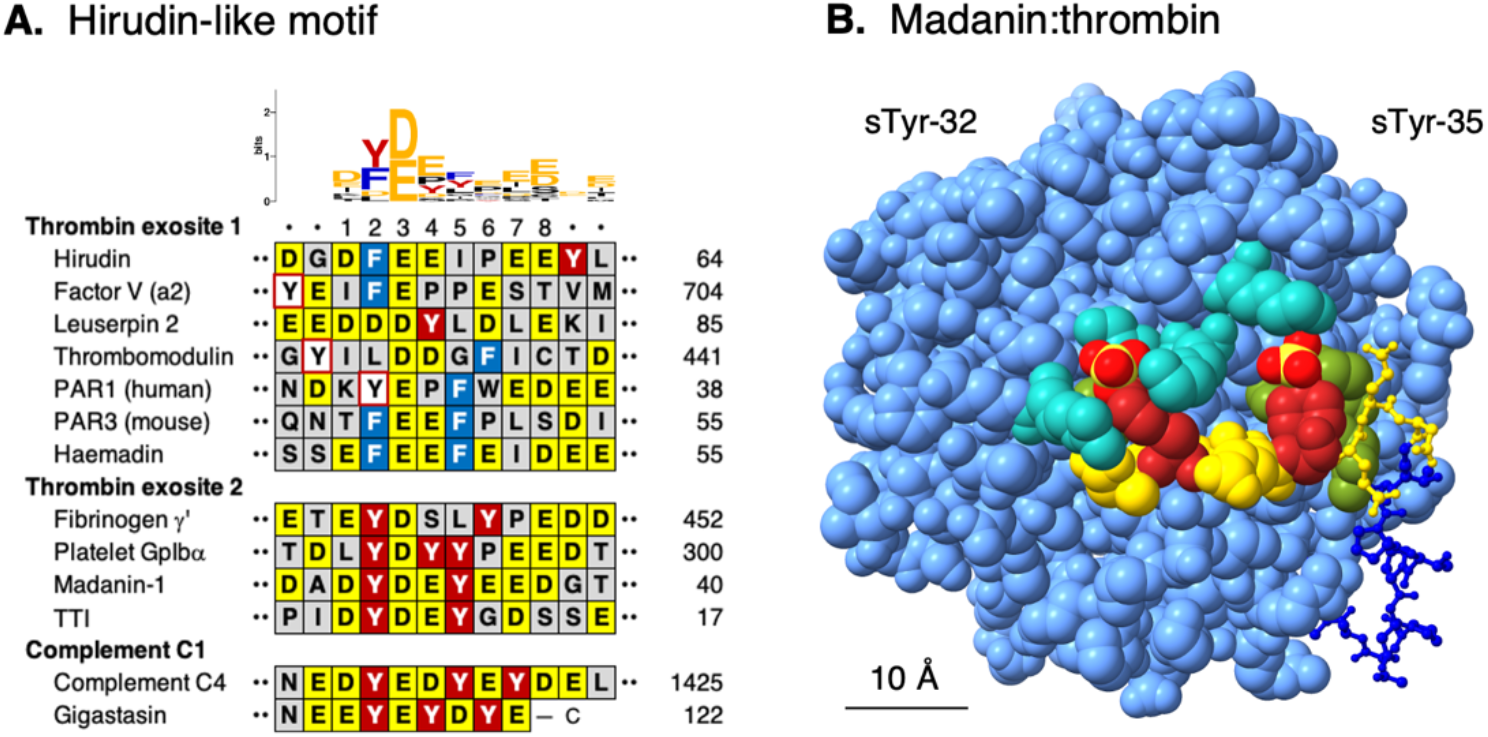
sTyr in hirudin-like motifs. one and one-half column figure (5” X 2.6”) **A**. sTyr cluster contexts are shown for thrombin-binding hirudin-like motifs from hirudin (PDB code 2PW8); factor V (a2 segment; 3P6Z); leuserpin 2 (1JMO); thrombomodulin (1DX5); peptide-activated receptor-1 (PAR1; human; 3LU9); peptide-activated receptor-3 (PAR3; mouse;2PUX); haemadin (1E0F); madanin-1 (5L6N); fibrinogen γ’ (2HWL); platelet GpIbα (4CH2); tsetse thrombin inhibitor (TTI; 6TKG); complement C4 (GenBank accession AAA51855); and gigastasin (5UBM). Residues are colored according to the key in **Fig. 3D**. The sequence logo (180,181) was generated from the alignment shown. The hirudin-like motif is defined here as positions 1-8. Positions 2 and 3 comprise a conserved aromatic-acidic residue pair, consisting of Phe, Tyr or sTyr at position 2 and Asp or Glu at position 3. The remaining hirudin-like motif residues are hydrophobic, negatively charged, or both in the case of sTyr, and binding to thrombin exosites involves both electrostatic and hydrophobic contacts (142). **B**. The co-crystal X-ray model of sulfated madanin bound to exosite 2 on thrombin (light blue) is depicted at 1.6 Å resolution (144) (PDB code 5L6N). Atoms are shown as Van der Waals spheres, except for madanin residues 36-60 shown as balls-and-sticks. Madanin residues sTyr-32 and sTyr-35 (dark red; S atoms yellow) make electrostatic interactions with exosite 2 Arg and Lys residues (light green) and hydrophobic interactions with other residues (olive green). Madanin Asp and Glu residues (yellow) are highlighted for reference.

Thrombin contains two cationic surface patches, termed anion-binding exosites, that bind different protein substrates through their hirudin-like motifs (137,138). Thrombin exosite 1 aligns both endoprotease substrates and DTIs such as hirudin with respect to the adjacent active site, whereas exosite 2 tethers thrombin at the site of blood clots, binding platelet surfaces via platelet **<u>g</u>**lyco**<u>p</u>**rotein **Ibα** (GpIbα) and to fibrin clots via fibrinogen γ’ (142,143). Exosite 2 also binds the DTIs madanin and **<u>t</u>**setse **<u>t</u>**hrombin **i**nhibitor (TTI) (51,144). Each exosite contains several basic Arg and Lys residues, enabling different contacts with acidic Asp, Glu and sTyr residues in different hirudin-like motif sequences.

As defined by the sequence alignments presented in **Fig. 5A**, the hirudin-like motif spans eight contiguous residues, of which most are acidic, aromatic, or both in the case of sTyr. Critically, hirudin-like motifs in all four proteins known to bind thrombin exosite 2 contain an sTyr cluster, whereas hirudin-like motifs from exosite 1-binding proteins, including hirudin itself, rarely contain sTyr residues and none at conserved positions (**Fig. 5A**).

Co-crystal X-ray structures of thrombin with exosite 2-bound sulfopeptides, illustrated here with the DTI madanin (144), reveal conserved contacts to thrombin residues across exosite 2 (24,51,145) (**Fig. 5B**). Residues sTyr-32^madanin^ and Asp-33^madanin^, which occupy the hirudin-like motif conserved positions 2 and 3, make both ionic and nonpolar contacts to numerous exosite 2 residues (144) that form the core interaction with thrombin (24). Residue sTyr-35^madanin^ is more exposed (**Fig. 5B**). Thrombin contacts to all three residues are conserved in all four available co-crystal X-ray structures (51), indicating that the sTyr cluster-containing hirudin-like motif provides a well-defined high-affinity binding determinant for exosite 2.

Peptide binding analyses with sulfopeptides (51,144) or phosphopeptides (24,145) illuminate these structural models, again illustrated here with sulfo-madanin. The thrombin inhibition constant for doubly sulfated madanin is about 400-fold lower than for non-sulfated madanin, showing the essentiality of sTyr. Madanin with only sTyr-35 is two-fold more effective than non-sulfated madanin, whereas madanin with only sTyr-32 is about 25-fold more effective (144). Thus, like chemokine receptor CCR5 interaction with gp120, the core sTyr residue (sTyr-32^madanin^; sTyr-14^CCR5^) is essential whereas the ancillary sTyr residue (sTyr-35^madanin^; sTyr-10^CCR5^) enhances binding only in the presence of the core (111,144) (**Fig. 5B**).

Like hemostasis, the complement system is activated through an endoprotease cascade (146). Complement component C1, a serine protease homologous to thrombin, has an anion-binding exosite that interacts with an sTyr cluster-containing hirudin-like motif in the substrate C4 (147,148) (**Fig. 5A**). The X-ray co-crystal structure of complement C1 bound to gigastasin, a C1 inhibitor made by *Haementaria* leeches, reveals electrostatic contacts from C1 exosite basic residues to sulfate O atoms from gigastasin residues sTyr-117 and sTyr-119 (149). Thus, the homologous hemostasis and complement systems share the use of hirudin-like motifs to direct partner protein binding to exosites.

Segments with sTyr clusters are prevalent in bloodstream activities, including leukocyte migration and signaling, blood clotting, complement activation and triglyceride metabolism. These activities also are modulated extensively by interactions with sulfated GAGs such as heparan, anionic oligosaccharides that help control hemostasis through binding thrombin (103,150-152). Indeed, thrombin exosite 2 was identified initially as the binding site for the highly-sulfated GAG heparin (142). Thus, a cluster of two or three sTyr residues can resemble a sulfated GAG to make conserved contacts with some of the many available basic residues in exosite 2, thereby expanding its valency beyond that of heparin-binding. Notably, basic exosite residues that contact GAG substrates (142) mostly are distinct from those that contact sTyr-containing substrates, illustrating the broad binding versatility provided by exosite 2.

### sTyr in acidic arrays

Finally, some sTyr residues lie within relatively long tracts of acidic Asp and Glu residues (**Fig. 2D**). Glycosyl**<u>p</u>**hosphatidyl**<u>i</u>**nositol-anchored **<u>h</u>**igh-density lipoprotein **<u>b</u>**inding **<u>p</u>**rotein **<u>1</u>** (GPIHBP1) is a membrane protein critical for triglyceride metabolism. The GPIHBP1 amino-terminal acidic tract requires a single sTyr residue for proper interaction and function with its binding partner, lipoprotein lipase (47). As with other examples above, the GPIHBP1 amino-terminal acidic tract is not visible in the lipoprotein lipase-GPIHBP1 co-crystal X-ray structure (153).

At the opposite extreme lie certain extracellular matrix proteins such as fibromodulin, whose amino terminus has an extended tract of acidic residues including several sTyr (154,155). This region binds collagen and a variety of heparin-binding proteins (156,157). The collagen-modifying enzyme lysyl oxidase has a similar sTyr-containing acidic tract through which it binds collagen (158). The fibrinogen and lysyl oxidase acidic tracts contain several Tyr residues, but in most cases, it has not been possible to determine which of these is sulfated to form sTyr (157,158).

### Challenges and prospects

The examples presented here exemplify the diverse contexts for the sTyr post-translational modification. A single sTyr residue can enhance binding affinity by several hundredfold. A cluster of two or three sTyr residues provides a wider range of binding modes, from versatile, as illustrated by the chemokine receptor CCR5 sTyr-containing amino-terminal segment, to stringent, as illustrated by hirudin-like motif interactions with thrombin exosite 2.

Post-translational modifications often work in concert, with different combinations exerting different effects (1). sTyr residues often occur with nearby *N*-or *O*-linked glycans, and differential glycosylation can affect function (2). For example, PSGL-1 residue Thr-16, near the sTyr cluster (**Fig. 4A**), is glycosylated in leukocytes to enable interaction with P-selectin, but it is not glycosylated in certain T cells wherein PSGL-1 interacts with VISTA (49,159). This encourages a parallel hypothesis, that differential tyrosyl sulfation also can help determine binding partner selection. Indeed, the chemokine receptor CXCR4 (for one example) requires residue sTyr-21 for chemokine ligand CXCL12 binding but not for HIV-1 entry (160). It is not known if or how CXCR4 tyrosine sulfation is regulated with respect to cell type or external stimulus.

The extent of sulfation at a given Tyr residue can be incomplete, such that only a fraction of proteins in a given population carry a particular sTyr residue (19,44). This is difficult to evaluate directly because MS analyses may result in sulfate loss (9). Does heterogeneous sulfation result from stochastic TPST catalysis? Or are different sulfation states (potentially with distinct ligand-binding properties) programmed in different cell types or in response to different stimuli?

Evaluating the location and extent of tyrosine sulfation also requires better understanding of how TPST engages its substrates and how its activity is regulated. Recently described TPST inhibitors may prove helpful (161). Substrates require nearby Asp or Glu residues for efficient sulfotransfer, but overall specificity determinants are not well-defined (70-72) in comparison to tyrosine kinases (162). However, animal cells make hundreds of tyrosine kinases but only two TPSTs. Therefore, TPST might catalyze some level of indiscriminate sulfation within a flexible anionic segment containing multiple Tyr residues. For example, several nearby Tyr residues within glycopeptide receptors and the C3a receptor are sulfated in vivo (**Fig. 3D**), even though sulfates at these positions are not required for receptor function (94,96).

Similarly, chemoreceptor CCR5 residue sTyr-3 (**Fig. 4A**) is considered to have minimal functional significance (109,110,163,164) and therefore usually is not studied (111,124). However, in vitro studies show that position Tyr-3 is the first to be sulfated in a CCR5 peptide that includes residue Met-1 (165,166). Does sTyr-3 result simply from “bystander” sulfation by TPST enzymes with broad substrate recognition? Or does it have a defined function (19)?

In vitro, pTyr can substitute for sTyr in some contexts (24,39,40) but not in others (164). Nevertheless, stringent specificity for sTyr in vivo is seen from the number of molecular mimics that contain sTyr, including anti-gp120 CD4i antibodies (115,117) as well as inhibitors of chemokine signaling (50), thrombin (26,51,144,167), and complement C1s (149). A striking example is the bacterial RaxX sulfopeptide, which acts as a molecular mimic of the plant PSY sulfopeptide hormone (76,77) (**Fig. 3A**). Although genes encoding TPST are ubiquitous in metazoa and plants, they are only sparsely distributed among relatively few bacterial lineages (168), and often are adjacent to genes encoding synthesis and export of extracytoplasmic proteins (V. Stewart and P. C. Ronald, unpublished observations). Thus, because RaxX requires sTyr for activity, the bacterium must synthesize TPST.

One potential research goal is to make defined sTyr residues to facilitate engineering of new interaction determinants. An initial proof-of-principle is eCD4-Ig, a chimeric protein that effectively blocks HIV-1 entry in rhesus macaques (118,169,170). eCD4-Ig contains an essential sTyr-rich peptide, derived from the sTyr-containing anti-gp120 antibody E51 variable region (**Fig. 4C**), and therefore is modeled closely after a pre-existing sTyr-containing segment.

More challenging will be to design sTyr interactions de novo. Placing sTyr residues at predetermined locations would require understanding and manipulating the substrate binding site as described above, or it could be accomplished through genetic code modification (29,32,33). Designing an effective sTyr binding site might be more feasible, given the wide range of naturally occurring sites characterized to date. An alternative approach is to use SH2 domains modified to recognize sTyr in place of pTyr (21).

A separate platform for engineering sTyr residues may come from **<u>ri</u>**bosomally synthesized and **<u>p</u>**ost-translationally modified **<u>p</u>**eptides (RiPPs), microbial natural products with diverse chemistry and potential applications (171). RiPPs are matured through a variety of post-translational modifications, and there is interest in engineering these modifications to create new RiPPs. The bacterial RaxST TPST, which synthesizes the sTyr residue in the RaxX RiPP, is a good candidate for strategies to introduce novel sTyr residues in engineered RiPPs (78,171). Additionally, certain polyketide biosynthesis complexes include SULT-type sulfotransferases among several modifying enzymes (172-174), so it may be possible to add TPST modules to non-ribosomal peptide synthesis complexes (175).

Finally, sTyr residues touch many topics in human medicine. For example, there are numerous sTyr residues in the homologous multidomain proteins, coagulation factors VIII (FVIII; six sTyr) and V (FV; seven sTyr) (176). Classic hemophilia results from FVIII deficiency, and one of the many causative *F8* alleles encodes a missense substitution of FVIII Phe for sTyr-1680 (176), documenting an important function for sTyr in hemostasis. Indeed, synthetic FVIII used for human therapy contains a full complement of sTyr residues (177). Nevertheless, relatively little is known about sTyr function in these critical hemostasis proteins (178,179). Research in FVIII and FV surely can benefit from the technical advances that now enable synthesis of sTyr residues at defined locations in vitro (25,27) and in vivo (29,32,33).

sTyr residues are versatile interaction determinants, essential for many critical interactions but also challenging to study. This means that sTyr residues are under-documented and -studied relative to other post-translational modifications.

Nevertheless, the accumulated knowledge base makes it easier to predict and analyze newly discovered sTyr residues, and such discovery will be facilitated as MS methods for sTyr detection increasingly are refined. One expects to see accelerated progress, not only in finding new sTyr residues but also in understanding functions for currently known examples.

## Supporting information

None.

## Acknowledgements

We thank Drs. Anna Joe, Dee Dee Luu and Chet Price for their helpful critiques on an early draft version of this manuscript, and the anonymous reviewers who provided candid, very helpful comments and suggestions.

## Author contributions

V. S. and P. C. R. wrote and edited the manuscript.

## Funding and additional information

Tyrosine sulfation research in the Ronald laboratory is supported by Public Health Service grant GM122968 from the National Institute of General Medical Sciences (awarded to P.C.R.). Molecular graphics and analyses were performed with UCSF Chimera, developed by the Resource for Biocomputing, Visualization, and Informatics at the University of California, San Francisco, with support from NIH P41-GM103311.

## Abbreviations

The abbreviations used are:

CCL: CC-type chemokine ligand
CCR: CC-type chemokine receptor
CD4i: CD4-induced
CIF: Casparian strip integrity factor
CXCR: CXC-type chemokine receptor
DTI: direct thrombin inhibitor
FSHR: follicle-stimulating hormone receptor
FV: coagulation factor
V; FVIII: coagulation factor
VIII; GAG: glycosaminoglycan
GpIbα: glycoprotein Ibα
gp120: glycoprotein spike 120 kDa subunit
HCD3: heavy-chain complementarity-determining region 3
LRR: leucine-rich repeat
PSY: plant peptide containing sulfated tyrosine
Rax: required for activation of Xa21-mediated immunity
RGF: root meristem growth factor
RiPP: ribosomally synthesized and post-translationally modified peptide
PSGL-1: P-selectin glycoprotein ligand-1
PAPS: 3’-phosphoadenosine 5’-phosphosulfate
PSK: phytosulfokine
SH2: *src* homology 2
sTyr: sulfotyrosyl residue
TPST: tyrosyl-protein sulfotransferase
TTI: tsetse thrombin inhibitor

Intentionally blank page to enable side-by-side figures and legends.

